# Quantification of Aquatic Unicellular Diazotrophs by Immunolabeled Flow Cytometry

**DOI:** 10.1101/2022.06.23.497322

**Authors:** Eyal Geisler, Hagar Siebner, Eyal Rahav, Edo Bar-Zeev

## Abstract

Quantifying the number of aquatic diazotrophs is highly challenging and relies mainly on microscopical approaches and/or molecular tools that are based on *nif* genes. However, it is still challenging to count diazotrophs, especially the unicellular fraction, despite their significant contribution to the aquatic nitrogen cycle. In this study a new method was developed to quantify unicellular diazotrophs by immunolabeling the nitrogenase enzyme followed by identification and quantification via flow cytometry. The new quantification method was initially developed using a diazotrophic monoculture (*Vibrio natriegens*) and verified by various auxiliary approaches. It was found that only 15-20% of the total number of *V. natriegens* cells have synthesized the nitrogenase enzyme, even though the media was anaerobic, and N limited. This approach was further tested in samples from marine and freshwater environments. It was found that the ratio of diazotrophs to total bacteria was 0.1% in the Mediterranean Sea, while 4.7% along the Jordan River. In contrast, the specific N_2_ fixation per unicellular diazotrophs was highest in the Mediterranean Sea (88 attomole N cell^-1^ d^-1^) while the total N_2_ fixation rates were lowest in the lake and the river (0.2 nmole N L^-1^ d^-1^). Overall, we expect that this direct quantification approach will provide new insights on the number and contribution of unicellular diazotrophs to total N_2_ fixation in marine and freshwater environments under various conditions.

## Introduction

Biological N_2_ fixation is a central process in marine and freshwater environments as it supplies new nitrogen compounds and support primary production (Gruber & Galloway, 2008; Zehr & Capone, 2020). Dinitrogen fixation is carried by a specific subgroup of bacteria and archaea known as diazotrophs. These organisms use the nitrogenase enzyme, a two-component complex comprised MoFe protein and Fe-reductase protein (Hoffman et al., 2014). Studies have indicated that aquatic diazotrophs include autotrophic (Zehr, 2011), heterotrophic (Bombar et al., 2016; Riemann et al., 2010) and mixotrophic (Benavides et al., 2020; Feng et al., 2010) metabolism to maintain the energetic requirements of the nitrogenase enzyme. Aquatic diazotrophs can be found in benthic mats, organized in long filamentous chains (up to few hundreds of micrometers), free living planktonic cells, or associated with aggregates (Bergman et al., 2013; Bertics et al., 2010; Riemann et al., 2022).

Quantifying the abundance of aquatic diazotrophs, especially the unicellular fraction, is challenging, thus missing in most ecological studies, despite their biochemical importance. Traditionally, microscopical approaches were used to count diazotrophs, for most part filamentous bacteria such as *Trichodesmium* sp. and *Anabena* sp. (Spungin et al., 2016; Zulkefli & Hwang, 2020). During the last two decades various molecular methods such as quantitative polymerase chain reaction were also used to estimate the numbers of smaller, unicellular diazotrophs (Foster et al., 2007; Turk et al., 2011). These methods are mostly based on evaluating the numbers of diazotrophs focus on the potential transcription (DNA) or expression (mRNA) of *nif*H genes. Previous studies have immunolabeled the nitrogenase enzyme to visualize aquatic diazotrophs (Currin et al., 1990; Geisler et al., 2019; Lin et al., 1998). Differently than targeting the *nif*H gene, it was estimated that diazotrophs which synthesized the nitrogenase enzyme are likely to actively fix dinitrogen. However, the advantages of nitrogenase immunolabeling for unicellular diazotrophs quantification was yet developed.

In this study nitrogenase immunolabeling was compiled with flow cytometry to quantify the number of unicellular diazotrophs in aquatic environments. The quantification method is based on immunolabeling the MoFe subunit of the nitrogenase enzyme by two antibodies conjugated to a green fluorophore. The method was first calibrated with *Vibrio natriegens* as a representative unicellular diazotrophs and then validated in-situ from several marine and freshwater environments. Complimentary N_2_ fixation measurements were undertaken, enabling to quantify the diazotrophic cell-specific activity.

## Materials and Methods

### Culturing unicellular diazotroph and non-diazotrophic bacteria

*Vibrio natriegens* (ATCC 14048) *and/or Escherichia coli* (ATCC 11303) were acclimated in a gas tight bottles (100 ml) containing Luria Bertani Broth media (LB, Merck Millipore, BD, 0083370) under anaerobic conditions (∼0.2 mg L^-1^ O_2_) at 26 °C overnight. The LB media used to grow the *V. natriegens* also included 1.5 % NaCl (Sigma Aldrich, 312525). Cells were further diluted to ∼5×10^6^ cells ml^-1^ and re-grown to a mid-logarithmic phase with an ∼2×10^8^ cells ml^-1^ for 1-2 h under the same conditions. Bacterial cells were centrifuged (3500 g for 6 min) to remove the LB and resuspended in artificial brackish water (1 ml) to a final cell concentration of ∼2×10^6^ cells ml^-1^. The chemical composition of the artificial media is detailed in the supporting information.

Triplicate biological replicates containing either *V. natriegens, E. coli* or both (1:1) were resuspended in N limited brackish water, enriched with ^15^N_2_ (99%, Cambridge Isotopes, lot #NLM-363-PK, final concentration 1% v:v). Monocultures or mixed cultures were then incubated for 48 hours under dark and anoxic conditions at 26 °C. Additional two bottles from each bacterial type were not enriched with ^15^N_2_ to determine their natural isotopes ratio. At the conclusion of the incubation, sub samples were analyzed for N_2_ fixation rates, colony forming units (CFU), as well as total bacterial abundance (BA) and diazotrophic abundance (DA) as detailed below. In addition, diazotrophs were visualized by capturing immunolabeled subsamples by confocal laser scanning microscopy (CLSM).

### Collection of natural diazotrophs

Surface waters were collected from three sampling locations, the South Eastern Mediterranean Sea, Qishon Estuary, and Sea of Galilee Lake (Table S1). Water were incubated in 1 L Nalgene bottles. N_2_ fixation rates were determined by enriching the samples with 15 % of dissolved ^15^N_2_ stock. Enriched samples were incubated for 48 h at room temperature under 12h light/dark conditions. Subsamples (1.7 ml) were collected at the end of the incubation for BA and DA analysis as well as diazotroph microlocalization.

### Analytical methods

#### Diazotrophs immunolabeling for flow cytometric analysis

Monoculture and natural samples (1.7 ml) were fixed with 50% glutaraldehyde (final concentration, 0.2 % Sigma-Aldrich, G7651), flash frozen in liquid nitrogen and stored at -80 °C until analyses. Samples were prepared by slow thawing at room temperature (Figure 1A). Next, ethylenediaminetetraacetic acid (EDTA, Sigma Aldrich, 03690) was added (final concentration of 5 µM) to chelate cations and facilitate aggregates dispersion (Bogler & Bar-Zeev, 2018). Samples were also sonicated in a bath sonicator for 6 min to disassociate cells from the aggregate matrix and one another. Subsamples (1 ml) were transferred and centrifuged for 10 minutes at x4000 g (a subsample was also collected for total bacterial abundance, detailed are provided below). The supernatant was cautiously discarded to maintain the bacterial pellet. Wash solution was prepared by mixing phosphate buffer saline (PBS) and Triton X-100 (T, final concentration of 0.1 %, Sigma Aldrich, X100), define hereafter as PBST. The wash solution was added to perforate the cell envelope. Samples were centrifuged for 10 minutes at x4000 g, while the supernatant was cautiously discarded. This washings-centrifugal cycle was repeated three times to increase the efficiency of cellular perforation. Fresh anti-nitrogenase antibody (3 µg ml^-1^, Agrisera Antibodies AS01 021A) was prepared with PBST and bovine albumin serum (BSA, filtered 0.2 µm, 1 mg ml^-1^, Sigma Aldrich A2153) to minimize unspecific antibody binding (Figure 1B). Samples were then incubated while slowly rotating (Benchmark Scientific Roto-Therm Plus, H2024) for one hour at room temperature to facilitate binding between the nitrogenase MoFe subunit and the primary antibody. Unbonded antibodies were removed by washing the samples three times with PBST similarly to the above. Washed samples were then incubated at room temperature in the dark for 45 min with the secondary antibody (3 µg ml^-1^, Thermo Fisher Scientific A-11039) conjugated to a green fluorophore (Alexa Fluor™ 488) with Ex spectra of 498 nm and Em of 520 nm (Figure 1C). Any untagged residues of the secondary antibodies were removed by washing the samples three times with PBST as described above. Immunolabeled samples were suspended with sterile PBS (1 ml) without any additions. Additionally, few controls were prepared following the above procedure and tested to evaluate the specific tagging of diazotrophs: (1) Negative control, namely PBST-BSA without any antibodies, to determine whether any autofluorescence could be detected; (2) no addition of the 1^st^ antibody (PBST-BSA with the 2^nd^ antibody only); and (3) no addition of the 2^nd^ antibody (PBST-BSA with the 1^st^ antibody only) to verify if any unspecific adsorption occurred. Additional control was to test unspecific tagging by applying the immunolabeling approach on non-diazotrophic (*E. coli*) bacteria. It should be noted that after each washing stage a subsample (100 µl) was collected to count the number of bacteria that were lost (Figure S1).

**Figure 1.**
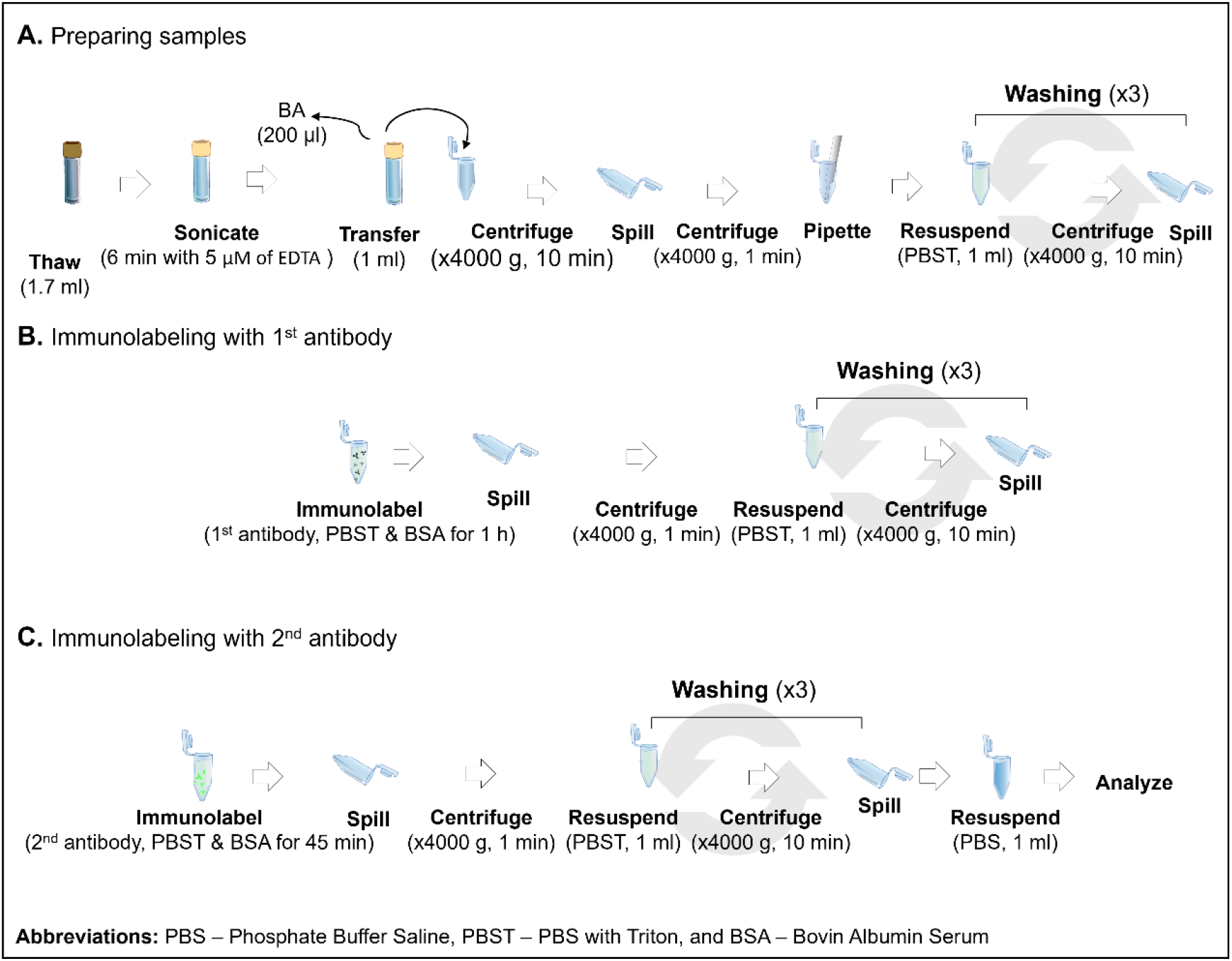
Graphical summary of the immunolabeling stage for flow cytometric analysis. The method is divided into three main stages: sample preparation (A); Immunolabeling with the 1^st^ antibody (B) and tagging with the 2^nd^ antibody conjugated to the green fluorophore (C).

### Counting immunolabeled diazotrophs and total bacteria using flow cytometry

Diazotroph abundance (DA) was determined by detecting and counting immunolabeled subsample (200 µl) using Attune-Next acoustic flow cytometer (Applied Biosystems). Changes in the abundance of monoculture diazotrophs were determined after diluting the immunolabeled samples (1:100, 1:250, 1:500, 1:1000). Monocultures were analyzed at a flow rate of 100 µl min^-1^, while reduced to 25 µl min^-1^ for natural samples. Stop condition was set to 20,000 counts for all samples. Calibration beads (1 µm, F8815, Invitrogen, Ex: 350 nm Em: 440 nm) were added (final concentration of 1.8×10^4^ beads ml^-1^) every 12 samples to evaluate the size spectrum of the sample.

Following the above, total bacterial abundance (BA) was quantified by staining non-immunolabeled subsamples (200 µl) with SYBR Green I (S7563, Invitrogen, final concentration, 1 nM) (Geisler et al., 2019). Samples were incubated for 15 min under dark conditions. Stained samples were measured with Attune-Next Acoustic Flow Cytometry. MilliQ (sterile) water samples were used to clean the system every five samples, while PBS were tested and subtracted as blanks. The specified lasers, dyes and filters are detailed in table 1.

**Table 1.**
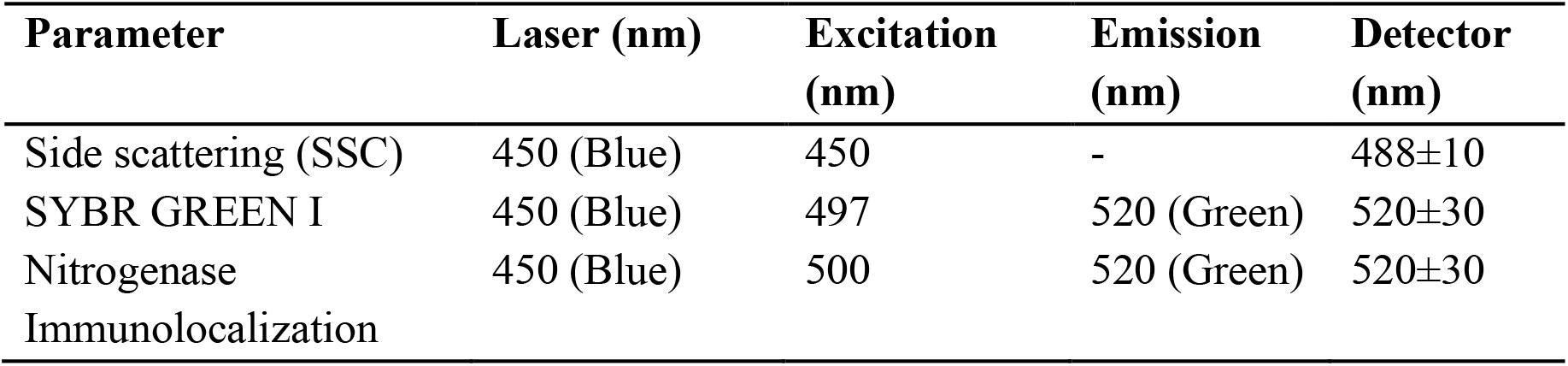
Analytical specifications of the flow cytometer.

| Parameter | Laser (nm) | Excitation (nm) | Emission (nm) | Detector (nm) |
| --- | --- | --- | --- | --- |
| Side scattering (SSC) | 450 (Blue) | 450 | - | 488±10 |
| SYBR GREEN I | 450 (Blue) | 497 | 520 (Green) | 520±30 |
| Nitrogenase Immunolocalization | 450 (Blue) | 500 | 520 (Green) | 520±30 |

### Quantifying cells using colony forming units (CFU) counts

Samples were collected from the incubation bottles and serial diluted in parallel to the immunolabeled samples (1:100, 1:250, 1:500, 1:1000). Subsamples (25 µl) were plated on an agar plate (1.5%, Bacto Agar, DF0140) with an N limited brackish water media (the recipe is detailed in the supporting information). CFU samples were incubated under anaerobic conditions for 48 hours at 26 °C. CFU were determined at the end of the incubation by counting plates with 30-300 colonies.

### Microlocalization of diazotrophs using confocal laser scanning microscopy (CLSM)

Immunolabeled samples (50 µl) were stained with 250 µg ml^-1^ of 40, 6-Diamidino-2-Phenylindole (DAPI, Ex 360 nm and Em 460 nm, Thermo Fisher, D1306) or (SYBR Green I, Ex 497 nm and Em 520 nm) and incubated for 20 minutes under dark conditions. Samples were drop-casted on a glass microscope slide, sealed by a cover slide and nail polish to minimize dehydration. Immunolabeled nitrogenase enzyme was visualized by a CLSM 900 equipped with a 488 nm laser (Power, 0.2 % digital gain, ∼500 V, pinhole, 32 µm). Stained bacteria were visualized with a 405 nm laser (Power, 0.9 %, digital gain, ∼730 V and pinhole, 37 µm). Samples were observed under a x63 lens (63x/1.4 Oil DIC M27, respectively). Non stained samples were used before to identify and subtract autofluorescence. Images were processed using Zeiss ZEN Blue edition (3.5, lite, Germany).

### Measuring N_2_ fixation rates

Samples were filtered on a pre-combusted glass microfiber filter (GF/F, Cytivia, 1825025, 450 °C, 4.5 h) after 48h incubation in ^15^N_2_ enriched media. Different volumes of samples (25 ml for lab cultures and 1L for environmental microcosms) were filtered, to ensure sufficient biomass (resulting in an amplitude of ∼1000 mv) on the filter as their source varied from monocultures to different natural environments. Samples were dried in the oven overnight (60 °C) and stored in the desiccator until measurements. Filters were carefully packed in tin capsules, with clean, pre-combusted glass fiber (GF/F) filters used as blanks. The samples were then analyzed using elemental analyzer (EA; Thermo Scientific, Flash 2000 HT) coupled with isotope ratio mass spectrophotometer (IRMS; Thermo Scientific, Delta V Plus). Working with filters and environmental samples over a wide range of concentrations requires caution during the isotopic measurement. The quality control measures taken during the measurement are detailed in the supporting information. Briefly: Three standards (Glutamic Acid USGS 40, Glycine USGS 64, and Caffeine USGS 62) were chosen for calibration, bracketing the expected range for δ^15^N of the enriched, as well as natural abundance samples, and ensure accuracy (Figure S2A). Acetanilide (Thermo Scientific, BN240741) was used for linearity test over the measured range (Figure S2B) and for quantitative calibration of peak amplitude vs. µg N in sample’s biomass (Figure S2D). A working range of 10 to 55 µg N per sample was determined to assure precision and avoid linearity effect. Similar ranges of nitrogen (> 10 µg N per filter) were previously determined (White et al., 2020). No drift was measured (slope = 0.01) and good precision was found (±0.3‰) throughout the analysis (Figure S2C). The natural abundance of ^15^N, reflected by the ratio of ^14^N/^15^N in each culture or environmental sample, was subtracted from that of the corresponding enriched sample to calculate N_2_ fixation rates, according to previous reports (Montoya et al., 1996).

### Statistical analyses

Statistical tests were ran using XLSTAT (2022.2, New-York). Before analyses, normal distribution of the data was validated using Shapiro Wilk test. The links between BA, DA, CFU and N_2_ fixation were measured by a Pearson correlation test. For comparing N_2_ fixation and BA/DA between samples, Analysis of Variance (ANOVA) was used with post-hoc Tukey test. All the tests were run under the confidence level of 95 % (α=0.05).

## Results and Discussion

### Detecting and quantifying a unicellular diazotrophic monoculture

Immunolabeled *V. natriegens* formed a distinct cluster after analyzing the samples by flow cytometry using a green detector over side scatter (Figure 2A). In contrast, only few unlabeled cells (< 1000 events) were captured in the same region (Figure 2B). Similarly, no *V. natriegens* cells were detected after tagging with the first or the second antibodies only (Figure 1C-D). Following the above, only conjunction of the two antibodies led to a positive detection of *V. natriegens* by flow cytometry, excluding any autofluorescence or unspecific adsorption of the tags to the cells. Finally, no immunolabeling by non-diazotroph, *E. coli* bacteria were detected in the region of interest by the flow cytometer (Figure 2E). The negative control highlighted that only cells with the nitrogenase enzyme could be tagged by the antibodies and detected as previously reported in other studies (Chelius & Triplett, 2000; Geisler et al., 2019).

**Figure 2.**
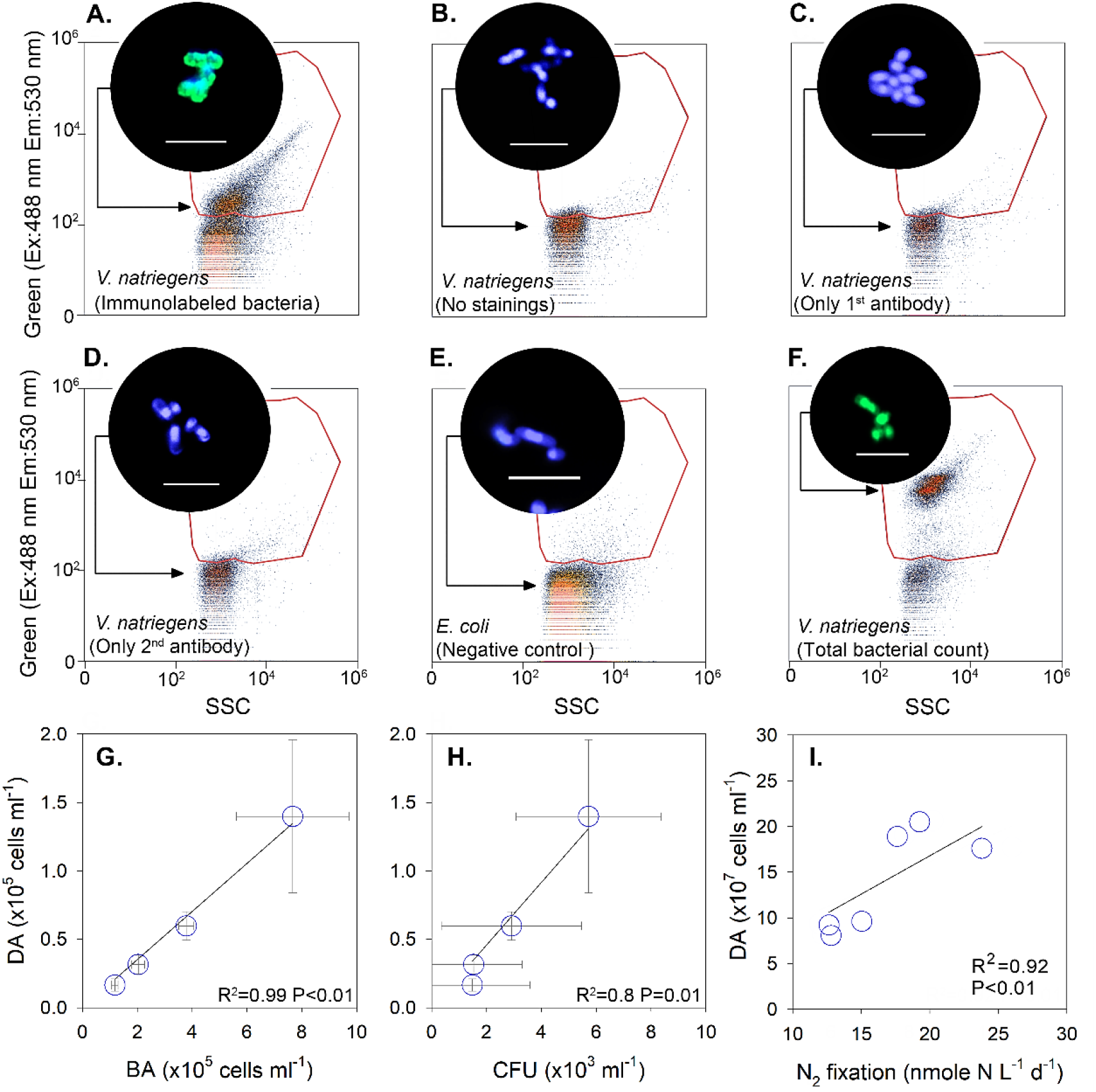
Representative density plots of the main verification tests (A-F) and correlation charts of the corresponding quantification results (G-I). The region of interest for immunolabeled *V. natriegens* diazotrophs as well as total bacterial count (individually tested after nucleic staining) was determined according to green fluorescence over side scatter (SSC). Diazotrophic abundance (DA) was correlated to total bacterial abundance (BA) (G) as well as colony forming units (CFU) (H) and N_2_ fixation rates (I). Top circles of each plot (A-F) capture subsample images using confocal laser scanning microscope with a scale bar of 5 µm. Diazotrophs were identified by nitrogenase immunolabeling (Green) while all the cells were detected by DAPI (Blue) or SYBR green (Green). Additional images are provided in supporting information (Figure S3A-E).

Total bacterial abundance was counted in an independent test after tagging a subsample with a nucleic acid stain (SYBR green) only. Tagging bacteria with SYBR green resulted in a distinct cluster that was identified in the same region of interest as described above (Figure 2F) and similar to previous studies (e.g., Geisler et al., 2019). Complimentary visualization of *V. natriegens* and *E. coli* subsamples by CLSM confirmed the results detected by the flow cytometry (Figure 2, circles).

Linear and significant correlation was detected between the number of immunolabeled *V. natriegens* and the total number of cells tagged by SYBR green from the same monoculture (Figure 2G). That trend line indicates that between 15 % to 20% of all *V. natriegens* bacteria were specifically tagged by immunolabeling, namely the cells that synthesized the nitrogenase enzyme. Correspondingly, a linear correlation was also found between immunolabeled cells and CFU counts that grew on limited nitrogen agar plates under anaerobic conditions for 48h (Figure 2H). It should be noted that the number of immunolabeled cells detected by flow cytometry was 20-25 times higher than those counted on the agar plates. In addition, a linear relationship was found between the number of *V. natriegens* that synthesized the nitrogenase enzyme and N_2_ fixation rates (Figure 2I), resulting in a specific N_2_ fixation per cell of 1.3±0.3 attomole N cell^-1^.

Lower percentage of free living diazotrophs that synthesized the nitrogenase enzyme compared to total cell count may indicate that heterotrophic N_2_ fixation was partly suppressed even under anerobic conditions and limited concentrations of inorganic nitrogen (confirmed also by low fixation rates per cell). Although the scope of the study was to develop a new quantification method for diazotrophs, it could be surmised that other constraints that were not measured such as pH (Luo et al., 2019) and/or carbon liability (Benavides et al., 2020; Rahav et al., 2016) impaired N_2_ fixation.

### Counting diazotrophic and non-diazotrophic mixed cultures

Two monocultures that included a diazotrophic (*V. natriegens*) and a non N_2_ fixing bacteria (*E. coli*) were mixed to test the differentiation capacity of the new immunolabeled—flow cytometry-based approach. Staining the DNA of subsamples with SYBR green for total bacterial count formed a distinct cluster (Figure 2F). Immunolabeling diazotrophic monoculture as well as a mixture of *V. natriegens* and *E. coli* bacteria resulted in a clear cluster over the conjugated nitrogenase (green) threshold (Figure 3A).

**Figure 3.**
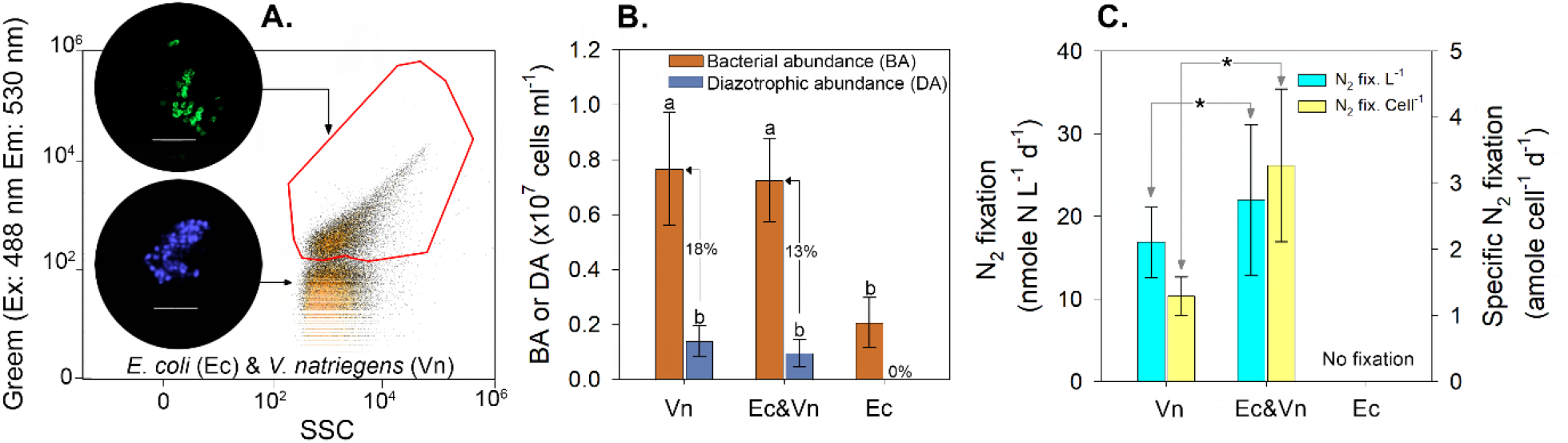
Representative flow cytometry plot of *E. coli* (Ec) and *V. natriegens* (Vn) mixed culture (A). The corresponding CLSM micrographs capture immunolabeled diazotrophs (5 µm scale bar). Additional image is provided in supporting information (Figure S3F). Abundance of non (*E. coli*) and diazotrophic bacteria (*V. natriegens*) in mono and mixed cultures was quantified after DNA staining and nitrogenase immunolabeling by flow cytometry (B). Corresponding N_2_ fixation rates were determined from all the cultures (C). N_2_ fixation rates per cell were calculated by normalizing the measured rates to the number of immunolabeled diazotrophs counted in the same culture. Values represent the mean and standard deviation from six independent replicates. Letters above the bars refer to ANOVA analysis followed by a Tukey post hoc test.

Quantifying total bacteria indicated that the numbers of *V. natriegens* only as well as a mixture with *E. coli* were similar (∼ 0.75 × 10^7^ cell ml^-1^) after 48 h of anaerobic incubation in a nitrogen limited media. It should be noted that the number of *E. coli* cells in a monoculture was lower by 71% (Figure 3B). Counting the immunolabeled cells indicated that the number of N_2_ fixing diazotrophs, namely *V. natriegens* that synthesized the nitrogenase enzyme, constitute 18% of the total *V. natriegens* cells and 13% of the mixed culture. Note, no immunolabeled *E. coli* cells were detected by the flow cytometer (Figure 3B), ruling out any unspecific links or adsorption of the fluorophore.

Corresponding N_2_ fixation rates were found to be significantly higher (1.3 times) by *V. natriegens* mixed with *E. coli* than in the monoculture (Figure 3C). That difference was even greater (2.5 times) when comparing N_2_ fixation rates per cell in the mixed culture to those measured from the *V. natriegens* culture. Altogether, it appears that mixing *V. natriegens* with a non diazotrophic heterotrophic bacteria such as *E. coli* spur N_2_ fixation rates per cell. It is plausible that increasing N_2_ fixation rates per cell enabled *V. natriegens* to compensate the enhanced consumption of limited dissolved inorganic nitrogen (initial concentration of 80 µM) that was included in the artificial media.

### Evaluating the abundance of unicellular diazotrophs in aquatic environments

Quantification of unicellular diazotrophs from different aquatic environments by immunolabeled flow cytometry resulted in a marked cluster, yet slightly more scattered than the monoculture controls (Figure 4A). Complimentary imaging of subsamples by CLSM indicated that only a small fraction of the cells collected from the Sea of Galilee Lake were tagged by nitrogenase immunolabeling (Figure 4A, top circle).

**Figure 4.**
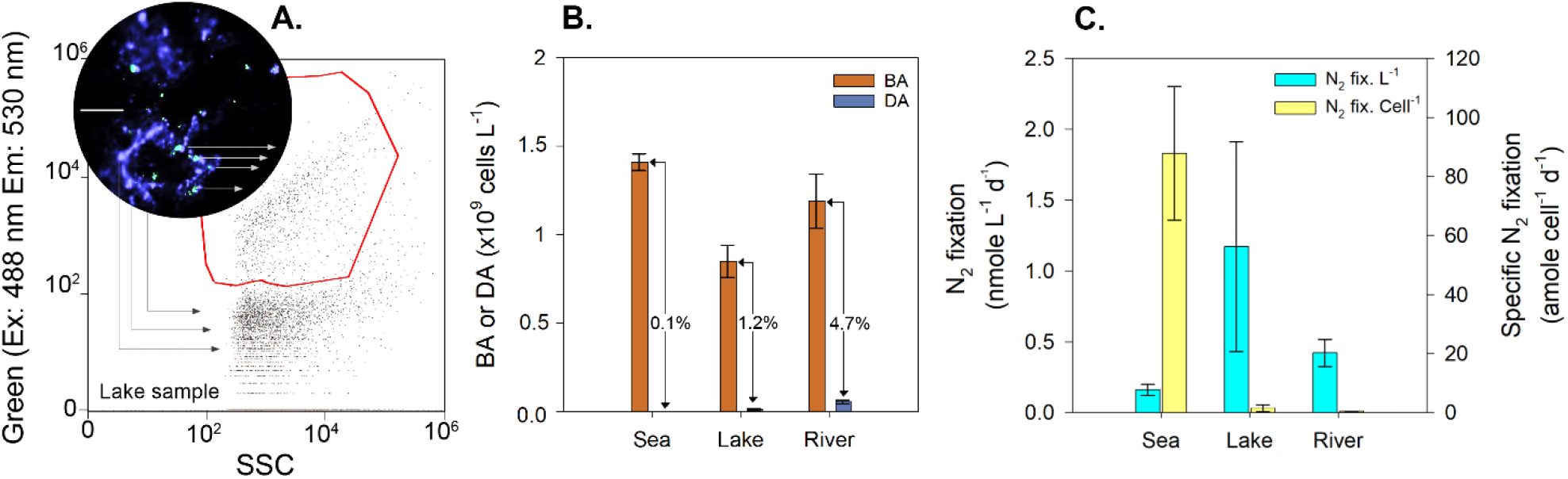
Representative density plots of an immunolabeled subsample (Green as a function of SSC) that was collected from the Sea of Galilee Lake (A). Complimentary CLSM image was further captured from a lake subsample (A, top circle), whereas diazotrophs were identified by nitrogenase immunolabeling (Green), while total bacteria were detected by DAPI (Blue). The image scale bar is 10 µm. Bacteria and diazotroph counts (BA and DA, respectively) were determined from the Sea of Galilee Lake, the Jordan River and the Mediterranean Sea (B). N_2_ fixation rates were measured from the same natural environments (C). N_2_ fixation per cell was calculated by normalizing the rates to number of DA.

Sporadic collection of water samples from different aquatic environments indicated that the abundance of unicellular diazotrophs ranged from 2±0.2 ×10^7^ cells L^-1^ in the Mediterranean Sea to 1±0.7 ×10^7^ cells L^-1^ in the Sea of Galilee Lake and 6±0.9 ×10^7^ cells L^-1^ in the Jordan River. The number of diazotrophs found in these environments were between 0.1% to 4.7% out of the total bacterial abundance (Figure 4B).

Corresponding N_2_ fixation rates from these samples were between 0.2 to 1.2 nmole N L^-1^ (Figure 4C), which are in similar ranges to previous reports (Halm et al., 2009; Marcarelli & Wurtsbaugh, 2006, 2009; Rahav et al., 2022). Note, data on N_2_ fixation rates in freshwater environments is still limited (Marcarelli et al., 2022). Normalizing these rates to the number of diazotrophs detected by immunolabeling flow cytometry resulted in N_2_ fixation per cells that ranged between 0.3-88 attomole N cell^-1^. These specific rates were found to be significantly lower at the Sea of Galilee Lake or the Jordan River than the Mediterranean Sea. Previous studies indicated on high specific rates correspond to high C:N or N:P ratios (Inomura et al., 2018; Knapp et al., 2012), conditions that are often found in the Mediterranean Sea. Differently, it was recently reported that N_2_ fixation rates per cell were high while at the same time nitrate concentrations in the surrounding environment were high (∼2 µM) (Mills et al., 2020), conditions that could potentially impair diazotrophy. Currently, the cellular mechanisms that control N_2_ fixation rates in environmental samples is not decisive and likely change according to the abiotic conditions and the different metabolic pathways.

## Conclusion

Coupling immunolabeling and flow cytometry can be used to quantify the number of unicellular diazotrophs that synthesized the nitrogenase enzyme, thus were likely fixing N_2_. This approach can be applied to count diazotrophs in controlled lab-scale experiments as well as various aquatic environments. Counting the total and immunolabeled cells of a diazotrophic monoculture indicated that even under anaerobic and N limiting conditions, only a fraction (15-20%) has synthesized the nitrogenase enzyme. That difference was likely due to the experimental conditions and yet highlights the importance of counting diazotrophs (even from monocultures under controlled conditions) to determine fundamental aspects such as specific N_2_ fixation per cell.

This approach can also enable quantification of unicellular diazotrophs in various marine and freshwater environments. Although the scope of this research was developing a new quantification approach for diazotrophs, it was interesting to find that N_2_ fixation per cell was highest in the oligotrophic Mediterranean Sea, compared to the Jordan River and the Sea of Galilee Lake, pointing on their potential significance for total biological N production.

It should be highlighted that this approach should be further investigated and developed: (i) differentiating and specifically counting unicellular heterotrophic or phototrophic diazotrophs is of high interest to estimate their contribution to total N_2_ fixation; (ii) the number of diazotrophs using this immunolabeled flow cytometry approach should be further compared to molecular-based approaches that are currently applied at various aquatic environments. Nevertheless, we suggest that adopting this approach could provide information on specific N_2_ fixation capacity of freshwater and marine diazotrophs. Moreover, quantifying diazotrophs will likely provide new insights on the contribution of these microorganisms to the aquatic nitrogen cycle.

## Acknowledgments

This paper is in partial fulfillment of EG Ph.D thesis at Ben Gurion University of the Negev. EB-Z thank the Israeli Science Foundation (grant number 944\21) for the supported of this project.

## Supporting information

A full recipe of brackish water is available for read. We also provided tests about washing efficiency of bacteria and quality control of EA-IRMS. Additional images of immunolabeled bacteria are also provided.

