## Supporting Information for "Quantification of Aquatic Unicellular Diazotrophs by Immunolabeled Flow Cytometry"

### Materials and methods

*Composition of the artificial brackish medium*: Sea salts (Advanced Pro Formula Salt, Royal Nature) were dissolved (15 g $L^{-1}$) in double-distilled water, filtrated through a coarse filter paper (Macherey-Nagel) and sterilized. Sodium bicarbonate was supplemented (15 mg $L^{-1}$) to provide additional buffering capacity while the pH was adjusted to 7–7.2. Vitamins (0.75 µM cobalamin, 4 µM biotin, and 0.8 µM thiamine HCl) and trace metals (4.8 mM $[FeCl_3]x6H_2O$, 17 µM $Na2EDTAx2H_2O$, 40 µM $[CuSO_4]x5H_2O$, 28 µM $[NaMoO_4]x2H_2O$, 76 µM $[ZnSO_4]x7H_2O$, 42 µM $[CoCl_2]x6H_2O$, and 1 mM $[MnCl_2]x4H_2O$) solutions were filtered through a 0.22-µm filter (Millex SLGV033RS) and added to the sterile media. Sterilized glucose (5 g $L^{-1}$) and $NH_4Cl$ (1.5 mg $L^{-1}$) solution was also added to the medium.

*Spatiotemporal characteristics of the samples collected from marine and freshwater environments.* Samples were collected from natural aquatic environments to test the newly developed immunolabeled approach and count unicellular diazotrophs from different marine and freshwater systems. The locations and sampling seasons were sporadically determined.

**Table 1:** Description of the different sampling locations and general abiotic conditions.

| Location | Lat. (N) | Lon. (E) | Salinity (ppt) | Water temp. (°C) | Trophic status |
|---|---|---|---|---|---|
| SE Mediterranean Sea | 34°30’02.5"N | 33°00’00.0"E | 39.5 | 18.3 | oligotrophic |
| Jordan River | 32°54’08.2”N | 35°36’50.0”E | ~0.09 | 17.2 | Mesotrophic |
| Sea of Galilee | 32°52’58.3”N | 35°36’34.4”E | ~0.25 | 19.2 | Mesotrophic |

*Bacterial loss during the immunolabeled approach.* One central concern was the present loss of bacteria during the different washing-centrifugation cycles. A significant loss of cells could result in underestimation of the corresponding counts. However, after optimization of the process as discussed in the main text (Figure 1), no significant change in the number of cells was found along the different washing cycles (Figure S1).

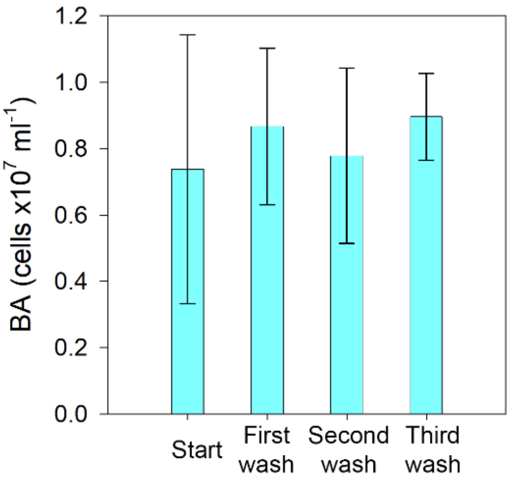


**Figure S1.** Bacterial abundance of *V. natriegens* along the immunolocalization process (n=two individual experiments). After every wash, sub samples (100 µl) were taken for counting bacteria. The staining process with SYBR GREEN I and analysis with a flow cytometry is done similarly as described in the main manuscript.

*Quality tests of EA-IRMS analyses*. The heterogeneity in biomass concentrations on the glass-fiber filters, which occasionally be found at low levels, requires great care in isotopic measurements. Assuring the accuracy and precision of the isotopic analyses was achieved by a set of quality control tests throughout the measurements. (a) Quartz crucibles were used and replaced every 25-30 samples to maintain the integrity and proper operation of the reactor. (b) Accuracy was determined by a calibration of the measured isotopic values with a set of secondary standards (Figure S2 A), bracketing the range of isotopic values of both natural abundance and enriched samples (USGS62 Caffeine, $\delta^{15}N_{AIR-N2}$=+20.17‰; USGHS64 Glycine, $\delta^{15}N_{AIR-N2}$=+1.76 ‰; and USGS40 L-Glutamic acid, -4.52 ± 0.06 ‰). (c) Linearity test was designed to verify the constancy of the isotopic value across a wide range of nitrogen levels in the samples and performed at the beginning of the work (Figure S2 B). According to the results, a lower detection limit of 10 µg N in samples was determined. This value is in accordance with previous publications (White et al., 2020). Standards were measured at the beginning and the end of each set. A representative standard (glycine USGS64) was measured repeatedly every 4-5 samples to verify the precision and correct for a potential drift when needed. No significant drift nor deviation was observed during the measurements (Figure S2C; slope = 0.01 $\delta^{15}N$ / µg N, standard deviation = 0.3). Finally, a calibration curve for the quantification of N amount in the biomass in samples by correlation to peak amplitude was plotted and verified using the different standards (Figure S2 D). A linear trend line was captured over a wide range of N masses, enabling reliable measurements of 10 to 55 µg N in samples.

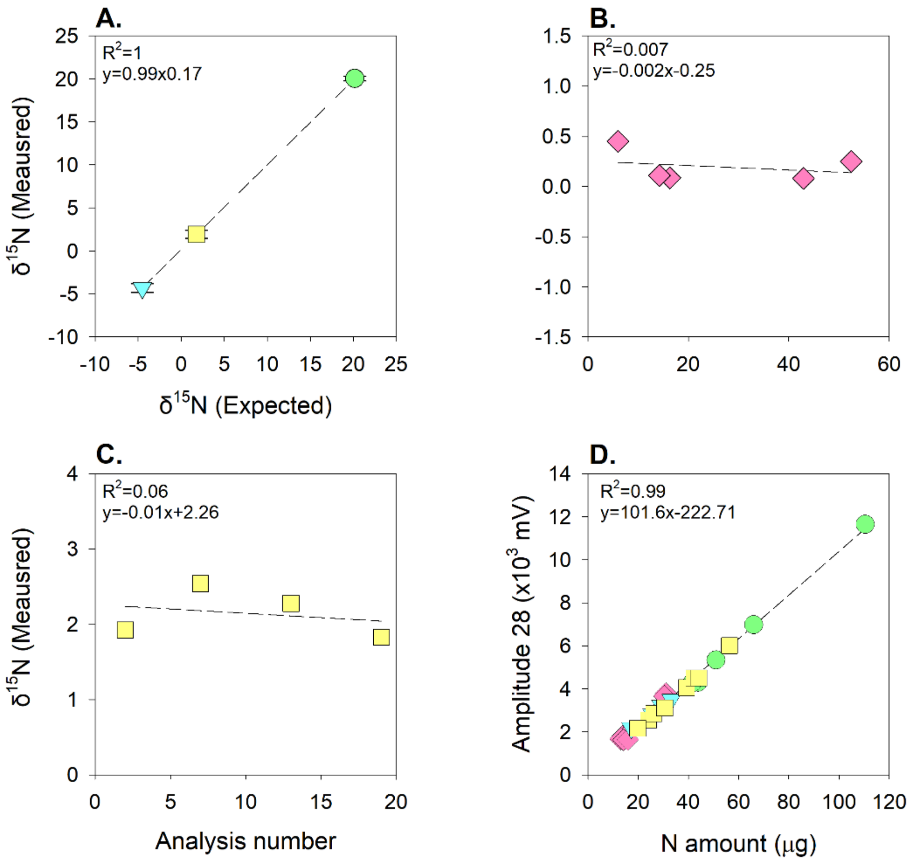


**Figure S2.** EA-IRMS quality control tests. Calibration curve was determined between expected and measured $\delta^{15}N$ for a range of known secondary standards (Caffeine USGS62, n=11; Glycine USGS64, n=18; Glutamic acid USGS40, n=11) (A). Linearity test of $\delta^{15}N$ in

Acetanilide over a range of N amount (μg) in samples (standard deviation for the range of 6 to 53 μg N was 0. 1) (B). Drift test of Glycine throughout the analysis (standard deviation was ±0.3 over 20 samples) (C). Calibration curve between N mass in the different standards to the measured amplitude, used for quantification the N in the sampled biomass (D). Glutamic acid (USGS40) was marked by blue triangles, Glycine (USGS64) by yellow squares, Caffeine (USGS62) by green circles, and Acetanilide by pink diamonds.

#### Results and discussion

Additional confocal laser scanning microscopy micrographs indicate the positive detection of immunolabeled nitrogenase using the two antibodies (*V. natriegens* Figure S3A) as well as the negative controls: no antibodies and absent of 1$^{st}$ or 2$^{nd}$ antibodies (Figure S3B-D). Additional samples were also included: negative control (non-diazotroph, *E. coli*) and mixed cultures (Figure 3S E-F).

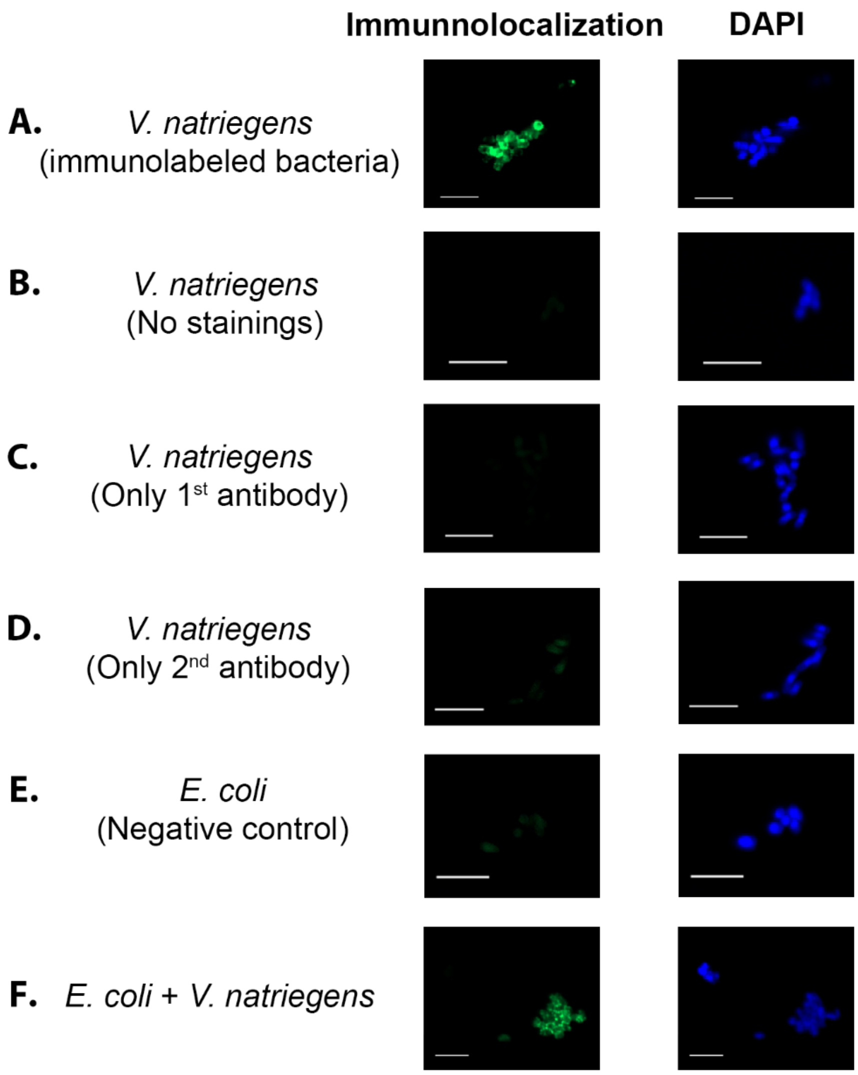


**Figure S3.** Immunolabeled micrographs of *V. natriegens* (diazotroph) and *E. coli* (non-diazotroph) bacteria. All samples have undergone the immunolabeled process (green) and DNA tagged by DAPI (blue). *V. natriegens* that were immunolabeled by both antigens were captured in green and positively tagged by DAPI (A). However, visualizing *V. natriegens* with no antibodies (B) or only one of them were only captured by DAPI (C,D). *E. coli* only was only found after DAPI staining (E). Mixture of *V. natriegens* (nitrogenase immunolabeled and DAPI tagged) and *E. coli* (stained only by DAPI) were visualized by the green and blue stains accordingly (F). The reported scale bar was 5 µm.
